# Comparative phylogenetic analysis of bacterial associates in Pyrrhocoroidea and evidence for ancient and persistent environmental symbiont reacquisition in Largidae (Hemiptera: Heteroptera)

**DOI:** 10.1101/064022

**Authors:** Eric Robert Lucien Gordon, Quinn McFrederick, Christiane Weirauch

**Author notes:** Corresponding author: Eric R. L. Gordon.

## Abstract

The ancient insect order Hemiptera, one of the most well-studied insect lineages with respect to bacterial symbioses, still contains major branches which lack robust phylogenies and comprehensive characterization of associated bacterial symbionts. The Pyrrhocoroidea (Largidae [220 species]; Pyrrhocoridae [~300 species]) is a superfamily of the primarily-herbivorous hemipteran infraorder Pentatomomorpha, though relationships to related superfamilies are controversial. Studies on bacterial symbionts of this group have focused on members of Pyrrhocoridae, but recent examination of species of two genera of Largidae demonstrated divergent symbiotic complexes between these putative sister families. We surveyed bacterial diversity of this group using paired-end Illumina and targeted Sanger sequencing of bacterial 16S amplicons of 30 pyrrhocoroid taxa, including 17 species of Largidae, in order to determine the identity of bacterial associates and similarity of associated microbial communities among species. We also constructed the first comprehensive phylogeny of this superfamily (4,800 bp; 5 loci; 57 ingroup + 12 outgroup taxa) in order accurately trace the evolution of symbiotic complexes among Pentatomomorpha. We undertook multiple lines of investigation (*i.e.*, experimental rearing, FISH microscopy, phylogenetic and co-evolutionary analyses) to understand potential transmission routes of largid symbionts. We found a prevalent, specific association of Largidae with plant-beneficial-environmental clade *Burkholderia* housed in midgut tubules. As in other distantly-related Heteroptera, symbiotic bacteria seem to be acquired from the environment every generation. We review current understanding of symbiotic complexes within the Pentatomomorpha and discuss means to further investigations of the evolution and function of these symbioses.

**Importance:** Obligate symbioses with bacteria are common in insects, particularly for Hemiptera wherein varied forms of symbiosis occur, though knowledge of symbionts remains incomplete for major lineages. Thus, an accurate understanding of how these partnerships evolved and changed over millions of years is not yet achievable. We contribute to our understanding of the evolution of symbiotic complexes in Hemiptera by characterizing bacterial associates of Pyrrhocoroidea focusing on the family Largidae and by constructing a phylogeny to establish evolutionary relationships of and within this group. Members of Largidae are associated with specific symbiotic *Burkholderia* from a different clade than *Burkholderia* symbionts in other Hemiptera and are members of the earliest-diverging superfamily of *Burkholderia*-associated Hemiptera. Evidence suggests that species of Largidae reacquire specific symbiotic bacteria every generation environmentally, a rare strategy for insects with potentially volatile evolutionary ramifications, but one that has persisted in Largidae and other related lineages since the Cretaceous.

## Introduction

With over 82,000 species, Hemiptera is one of the most diverse lineages of animals and is the most speciose order of insects which do not undergo complete metamorphosis (1). Their success can be tied directly to bacterial symbionts, which presumably allowed the ancestor of this lineage to exploit nutrient-limited diets of plant tissues such as xylem and phloem (2). Three lineages (Sternorrhyncha, Auchenorrhyncha, Coleorrhyncha) are exclusively herbivorous and nearly all constituent species remain associated with obligate intracellular symbionts that are vertically transmitted to offspring before oviposition (3–5). These obligate partnerships with bacteria may have helped to promote speciation through rapid development of potential genetic incompatibilities of symbiont and hosts resulting in higher rates of hybrid mortality (6). Sources of incompatibilities and other potentially deleterious mutations inevitably occur during population bottlenecks of intracellular symbiont lineages during transmission between generations and can eventually lead to compensatory mutations in host genomes (6). This comparatively rapid evolution of both symbiont and host can effectively cripple the symbiosis over many generations (7, 8). In many lineages, symbiont genomes become so reduced that former metabolic capability has been supplanted by secondarily-acquired symbionts or partitioned by speciation of one endosymbiont into two metabolically-distinct species (9–11).

Unlike other Hemiptera, the ancestor of the Heteroptera achieved independence from obligate symbionts at some point while transitioning from an herbivore to a predator (evolution of trophic strategies of Heteroptera summarized in [12]). However, two diverse clades of Heteroptera have secondarily re-evolved herbivory and together comprise more than 60% of heteropteran diversity, including many economically important pests of crops (13). In one of these radiations, Trichophora (Pentatomomorpha excluding Aradoidea), most members possess large populations of particular groups of extracellular symbiotic bacteria in blind tubules of the posterior midgut called caeca (14, 15). In the Pentatomoidea, these bacteria comprise various lineages of gammaproteobacteria which are primarily vertically transmitted via egg smearing, coprophagy or co-deposition in a jelly or capsule (16–19). Certain lineages of Lygaeoidea do not possess caeca and instead are associated with various bacteriome-inhabiting, vertically-transmitted gammaproteobacteria (20, 21), but all examined caeca-possessing members of the superfamilies Lygaeoidea (Berytidae, Blissidae in part, Rhyparochromidae, Pachygronthidae) and Coreoidea (Coreidae, Alydidae) are symbiotically partnered with members of *Burkholderia* (22). The mechanism of inoculation is from the environment (23–25) with young instar nymphs acquiring particular symbionts from soil (26), although some vertical transmission occurs in at least some lineages, such as Blissidae (27, 28). Caeca-possessing host species possess a symbiont-sorting organ at the junction of the third and fourth midgut section which blocks the passage of food and allows selective passage of particular bacteria (29).

In comparison to other hemipterans, herbivorous heteropterans tend to feed on relatively nutrient-rich parts of plants *e.g.* seeds, fruits or new buds. Despite this comparatively rich diet, hosts tend to face moderate to severe fitness deficits when deprived of beneficial symbionts (30–32). In Pentatomoidea, the function of symbionts is likely supplementation of amino acids as has been demonstrated for Urostylididae (19) and suspected for Pentatomidae and Plastapidae, where symbionts with reduced genomes retain capabilities to produce amino acids (33, 34). In Parastrachiidae (Pentatomoidea), symbionts recycle uric acid during long periods of diapause between availability of berries of their host plant (35). In Alydidae (Coreoidea), gene knockout experiments have demonstrated some requirements for effective symbiotic colonization, such as flagellar motility (29), and biosynthetic genes responsible for cell walls (36), secondary messenger, c-di-GMP (37, 38), and a bacterial energy storage polymer (39). These symbionts have been shown to enhance host innate immunity (40) or confer resistance to fenitrothion (41) but the primary function of *Burkholderia* symbionts to hosts remains unknown.

The Largidae together with Pyrrhocoridae, constitute the superfamily Pyrrhocoroidea which contains ~520 extant species (42, 43). Members of Pyrrhocoridae are associated with a characteristic microbiota primarily in the third section of the midgut which includes two obligate actinobacterial species, *Coriobacterium glomerans* and *Gordonibacter* sp. passed on via smearing of eggs (32, 44, 45). Only female Pyrrhocoridae are known to possess caeca though they appear to be vestigial and do not contain bacteria (45). Elimination of symbionts by egg sterilization results in host mortality and symbionts are thought to help their specialist hosts subsist on seeds of Malvales through supplementation of B vitamins (46). In contrast to Pyrrhocoridae, most largids appear to be generalist seed-feeding herbivores although some Old World genera may be associated preferentially with seeds of Euphorbiaceae (13, 47, 48). Early microscopic studies have demonstrated bacteria within caeca of members of Largidae previously identified as *Bacillus* or *Lactobacillus (14, 49, 50)*. While conducting a survey of bacterial diversity of Pyrrhocoridae, Sudarakaran et al. also discovered that representatives of the largid genera, *Largus* and *Physopelta*, are associated with *Burkholderia (51)*. A focused study on members of *Physopelta* found *Burkholderia* in midgut caeca from the plant-associated beneficial and environmental (PBE) clade (52). All other known heteropteran *Burkholderia* symbionts belong to a clade called the stinkbug-associated and beneficial environmental (SBE) clade with the exception of one family, Blissidae, in which *Burkholderia* isolates may belong to the former two clades as well as a third *Burkholderia cepacia* complex (BCC) clade. (27, 28).

Alternate scenarios of the evolution of bacterial symbioses within this group of herbivorous true bugs may be invoked depending on the phylogenetic relationship of Pyrrhocoroidea with respect to related superfamilies. Evolutionary relationships of this superfamily have been controversial, with recent molecular phylogenies (listed in chronological order) placing Pyrrhocoroidea as sister to Coreoidea + Lygaeoidea (53), Alydidae (54), Coreoidea (55) or Lygaeoidea (56) and the most recent morphology-based cladistic analysis finding this superfamily as sister to Coreoidea (57). The monophyly of Largidae has also been questioned proposing that Pyrrhocoridae may have evolved from Largidae (58), or specifically, that the old World tribe Physopeltini may be more closely related to Pyrrhocoridae than to the New World Largini (43). Establishing the relationships of this superfamily will allow for a more accurate explanation of how different symbiotic complexes evolved among and within this group, and allow for testing concordance among host and symbiont phylogenies.

In the current study, we aim to identify symbionts across Pyrrhocoroidea with a focus on the family Largidae and determine the relations of these symbionts to other known heteropteran symbionts through an Illumina bacterial 16S amplicon survey and targeted full-length 16s rRNA gene sequencing. We also construct the first comprehensive phylogeny of Pyrrhocoroidea to accurately determine the evolutionary history of this group and how this branch on the tree of life relates to others with bacterial symbionts. We seek evidence to determine the method of transmission of Largidae symbionts to offspring through experimental rearing, fluorescence in situ hybridization (FISH) microscopy, culturing and investigation of patterns of symbiont and host phylogenies. We summarize our results and the current knowledge on evolution of bacterial symbiont complexes in pentatomomorphan Heteroptera.

## Methods

### Rearing of *Largus californicus*

Adult specimens were captured from Lytle Creek in San Bernardino National Forest, San Bernardino County, CA in June 2015 and were enclosed in plastic containers with soil substrate from UCR campus (not from field environment) and held at room temperature. Specimens were also provided with grapes, cabbage and water in a vial plugged with cotton. An egg batch (of approximately 70 eggs) was divided and approximately half of the eggs were washed with distilled water and DNA was extracted from washed eggs and wash with a QIAGEN DNeasy Blood and Tissue kit. The other half of the eggs were allowed to hatch, which they did after two weeks (on July 28th, 2015) and five first instar nymphs were pooled together and subjected to DNA extraction two days after hatching. The caeca-containing region of the midgut of an adult specimen was also dissected and DNA extracted. The presence of bacteria in DNA extracts was analyzed with PCR with universal and genus-specific bacterial 16S primers (Table S1).

### Culturing

The caeca-containing region of the midgut of one adult specimen of *L. californicus* was dissected away from other gut tissue and macerated with an Eppendorf pestle for 2 minutes in 200 ul of PBS buffer (pH 7.4). An inoculating loop was used to spread the resulting cloudy homogenate liquid on a Luria-Bertani agar plate and incubated for three days at 37°C. Several representatives of dominant colony morphotype were analyzed using colony PCR with universal bacterial primers (Table S2) and the resulting sequences were blasted against GenBank after cleaning and sequencing of PCR products to confirm the identity of the bacterial isolate.

### Florescence in situ hybridization (FISH)

Gut tissue from live specimens of *Largus californicus*, after anesthetization at −20°C for three minutes, was dissected and stored separately in acetone as well as ~10 whole eggs from the unwashed half of the egg batch for whole mount microscopic preparations. We followed the protocol in (59), fixing tissue with Carnoy's solution overnight and staining gut tissue with DAPI for labelingl DNA, and two oligonucleotides probes: Cy-5-labeled universal bacterial probe (EUB-338; [60]) and Cy-3-labeled *Burkholderia*-specific 16S oligonucleotide (Burk129; [22]) from biomers.net for Cy-5 and Integrated DNA Technologies for Cy-3 (with HPLC purification) for staining of specific bacterial symbionts. Confocal microscopy was conducted with a Leica TCS SP5 using 405, 543 and 655 nm lasers for visualization of DAPI, Cy-3 and Cy-5 respectively.

### Sampling and DNA extraction

Individual specimens of all available of Pyrrhocoroidea (from the worldwide-ethanol collection of Heteroptera of the Weirauch lab; details listed in Table S1) along with two outgroup taxa (32 species total including 13 genera of Pyrrhocoroidea) were surface sterilized with a 1% bleach solution for 2 minutes and rinsed with 100% ethanol before removal of the abdomen from the thorax. Internal abdominal tissue was removed with sterile forceps and the resulting material was homogenized with a bead beater for 3 minutes at 30 Hz after addition of 100 μL of 0.1 mm glass beads and one 2.38 mm metal bead. Forceps were washed with EtOH, flamed and sterilized with a 10% bleach solution before and after each extraction. Each sample was incubated at 55° for 24 hours after addition of 10 μl of 800 U/ml Proteinase K before proceeding with DNA extraction.

### Host phylogeny

A total of ~4,800 bp of host DNA consisting of two mitochondrial protein-encoding (COI and COII) and three ribosomal genes (16S, 18S, 28S) were amplified from DNA extracts with primers listed in Table S2. PCR products were cleaned with Bioline Sureclean and submitted to Macrogen for Sanger sequencing. Chromatographs were edited in Sequencher v4.8 and aligned with MAFFT (E-INS-i strategy). A RAxML maximum likelihood phylogeny was constructed after partitioning based on gene and codon position for protein encoding genes and designation of the two included taxa of Pentatomoidea as an outgroup. Relevant sequences from GenBank were included for a maximally comprehensive phylogeny of Pyrrhocoroidea (57 taxa and at least 20 genera) and all newly acquired sequences (accessions numbers KX523359-KX523485) have been deposited on GenBank (Table S1). We noted that published sequences from *Dindymus lanius* on GenBank clustered closely with *Antilochus* in contrast to our own sequences from three species of *Dindymus* which included *Dindymus lanius* so we excluded the former sequences from our analysis in case of possible misidentification. All sampled representatives have been imaged with a Leica Microsystems imaging system and databased using Arthropod Easy Capture (AEC) implemented in the Plant Bug PBI and these data are publically available via “Heteroptera Species Pages” (http://research.amnh.org/pbi/heteropteraspeciespage).

### Illumina 16S amplicons sequencing and analysis

Gut tissue from a subset of 17 taxa was subject to Illumina amplicon sequencing of a ~300 bp fragment of the bacterial 16S gene using 799F and 1115R primers with barcodes for multiplexing, as in (61), intended to minimize amplification of chloroplast DNA (62) and conducted in triplicate PCR (35 cycles annealing at 52°) with 5 PRIME HotMasterMix. Triplicate PCR products were pooled and cleaned with Ultraclean PCR cleanup kit (MoBio, Carlsbad, CA). Illumina adaptors were added to templates via PCR on 1 μl of cleaned product with HPLC purified primers (15 cycles annealing at 58°; Table S2). Eighteen microliters of PCR product for each sample was normalized with a 96-well SequalPrep™ Normalization Plate and 5 μl of each normalized sample was pooled and assessed for quality with a 2100 Bioanalyser (Agilent, Santa Clara, CA) and the University of California, Riverside Institute for Integrative Genome Biology.

Samples were multiplexed on an Illumina Miseq lane with a MiSeq Reagent Kit v3 with 2 x 300 paired-end sequencing. Paired-end read was assembled, trimmed (reads with ambiguous bases or aberrant lengths removed) and demultiplexed using mothur v1.35.1 (63). Sequences were aligned to the Silva v4 reference alignment and checked for chimeras with the UCHIME algorithm using the most abundant sequences as a reference and assigned to OTUs at a 97% identity level and OTUs were classified using a Bayesian classifier both implemented in mothur (63). Jaccard and Bray Curtis community dissimilarity metrics along with weighted and unweighted Unifrac distance matrices (64) were computed after rarefying dataset to 2,066 reads per sample (the lowest number of reads in any sample after filtering) and clustered via PCoA ordination plots visualized with Plotly (65). Raw data are available on the NCBI Sequence Read Archive (SRA) under accession number SRP078165.

For visualization of OTU abundance with a heat map, we removed all OTUs which together comprised <1% of the total dataset and any representatives of any OTU which constituted less than 0.5% of the total reads in a sample. Together, these represent slightly less than 2.5% of total reads after trimming. The most common representative of any OTU unclassified at a genus level was blasted against GenBank and manually curated to the lowest level possible based on highest scoring blastn to known organisms. Putative chimeric OTUs based on high blast hit identity to distantly related bacterial lineages were removed. OTUs with identical top blast hits were combined for visualization purposes. Read abundance was plotted on the log scale after division by 10 (equivalent to log_10_ (reads) - 1) and plotted with the ggplot2 package in R.

### Targeted full-length 16S PCR and phylogeny

Full length 16S sequences were retrieved from two transcriptomes of *Largus californicus* sequenced recently as a part of the Hemipteroid Tree of Life project. Based on these sequences, we modified existing primers to amplify full 16S sequences of *Burkholderia* with two sets of PCR with combination of PBE-specific primers and *Burkholderia*-specific primers (Primers in Table S2; newly acquired 16S sequences KX527603-KX527621 in Table S1). PCR products were cleaned with Bioline SureClean and sequences were processed as for host genes. A comprehensive set of 272 ribosomal 16S sequences for environmental and insect associated isolates of *Burkholderia* along with named *Burkholderia* species and 3 *Pandoraea* outgroups was downloaded from GenBank and a tree for the genus including full length 16S sequences of dominant *Burkholderia* obtained from Largidae samples via PCR and representative *Burkholderia* reads (300 bp) of less dominant OTUs from the Illumina dataset chosen from each sample in which they were present was constructed (Fig. S2). Taxa were pruned from this tree using Mesquite v3.04 (66) for easier visualization, retaining close relatives of newly sequenced bacterial 16S genes, representatives of other insect-associated lineages and named species and the resulting dataset was realigned and phylogeny reconstructed (Fig. 3).

A cophylogenetic analysis of *Burkholderia* and host phylogenies was performed with TreeMap 3 (67) and Parafit (68). A maximally reconciled set branching pattern of host and symbiont phylogenies was visualized with TreeMap 3. Patristic distances were calculated with the cophenetic function in the ape package in R and the resulting phylogenetic relationships were tested for correlation of bacterial and host phylogenies via the statistical test implemented in Parafit with 10,000 permutations.

## Results

### Gut morphology, rearing, culturing

As has been previously described for congeners (Fig. 1B inset), the gut morphology of *Largus californicus* (adult pictured Fig. 1A) consists of five morphologically distinct midgut sections (Fig. 1B). The third section is the largest and most voluminous and is followed by a constricted region (Fig. 1G), homologous to the symbiont sorting organs described in other Heteroptera (29). Caeca are relatively short and numerous in the fifth midgut section consisting of two rows of tubes (Fig. 1B,C,F) and tend to be closely associated with the distal part of the Malpighian tubules *in situ*. FISH microscopy highlights a large density of bacteria in the caeca (Fig. 1D), which were stained with a genus-specific probe for *Burkholderia* (Fig. 1E). Egg batches did not contain any prominent co-deposited substance (Fig. 1H), however first instar nymphs did probe remains of hatched eggs (Fig. 1I, J).

**Figure 1.**
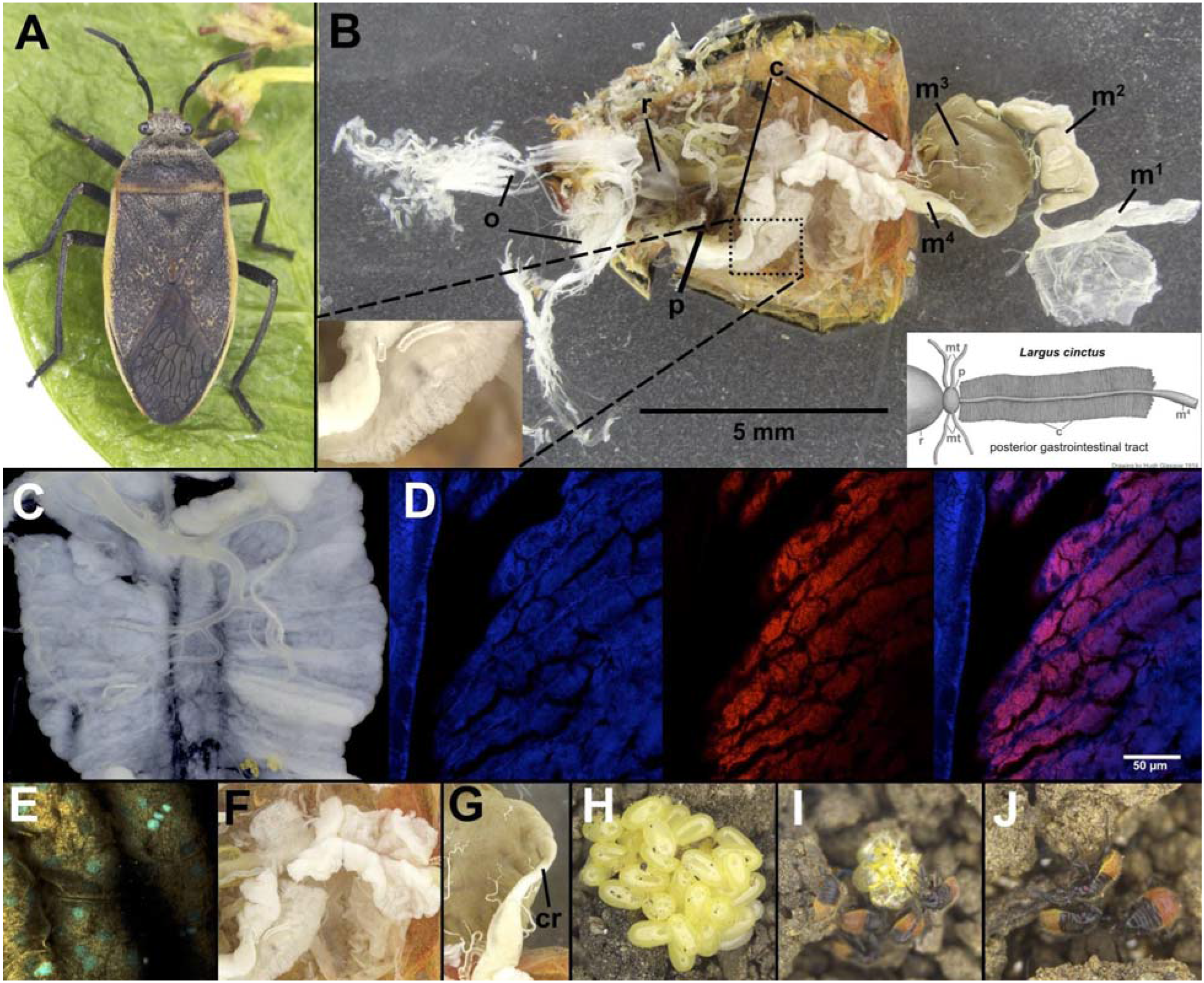
**A.** *Largus californicus* adult female. **B.** Gut morphology of *L. californicus*; Inset: Magnified posterior midgut caeca (left) and drawing of *Largus cinctus* posterior gastrointestinal tract (modified from Glasgow 1914; right). **C.** Caeca dissected away from other parts of gastrointestinal tract. **D.** FISH micrographs of caeca stained with DAPI (blue) and Cy-5 universal bacterial probe (red) for 16S; Left: 405 nm laser; Middle: 655 nm laser; Right: Merged. **E.** Merged FISH micrograph of caecal tissue stained with DAPI-only (blue) or with DAPI and Cy-3 *Burkholderia*-specific probe (orange). **F.** Magnified view of caeca-containing section of midgut. Anterior of gut oriented towards right. **G.** Close up view of constricted region between fourth and fifth midgut region. Anterior of gut oriented towards top. **H.** Egg batch after removal of about half of eggs. **I.** 1st instar nymph of *L. californicus* probing egg batch with labium. **J.** 1st instar nymphs of *L. californicus*.

Sequencing of PCR products amplified with general bacterial primers (Table S2) on DNA extracts of the isolated caeca-containing region (Fig. 1C) produced a clean chromatograph sequence matching *Burkholderia* (Table S1). DNA extracts of unwashed and washed eggs, first instar nymphs and the wash from washed eggs produced no product when assayed with PCR with *Burkholderia*-specific primers. Both sets of eggs and first instar nymphs did produce a band when assayed with general bacterial primers which when sequenced produced a clean chromatograph matching *Rickettsia* (Table S1). The *Burkholderia* symbiont was apparently easily cultured on LB plates as all sequenced representatives of the dominant small yellow colonies after plating of homogenized caeca were identical in sequence to each other and 100% identical to the PBE-clade *Burkholderia australis*, isolated from sugarcane roots, JQ994113.1 [69]).

### Phylogenetic relationships within and among Pyrrhocoroidea

Our phylogeny reconstructed Pyrrhocoroidea as sister to Coreoidea + Lygaeoidea (Fig. 2; Fig S1) with a bootstrap support value >85%. We recover, with moderate support, a monophyletic Largidae with a sister group relationship between the two subfamilies Physopeltinae and Larginae. The two predatory genera, *Antilochus* which are specialist predators of other Pyrrhocoridae (70), and *Dindymus* which possess variable trophic strategies though some are specialist predators of mollusks (71), do not appear to be most closely related to each other. In the unidentified female pyrrhocorid specimen L54, which may belong to the genus *Ectatops* or *Saldoides*, we observed caeca when dissecting gut tissue that were well-developed and comparable to those seen in *Largus*.

**Figure 2.**
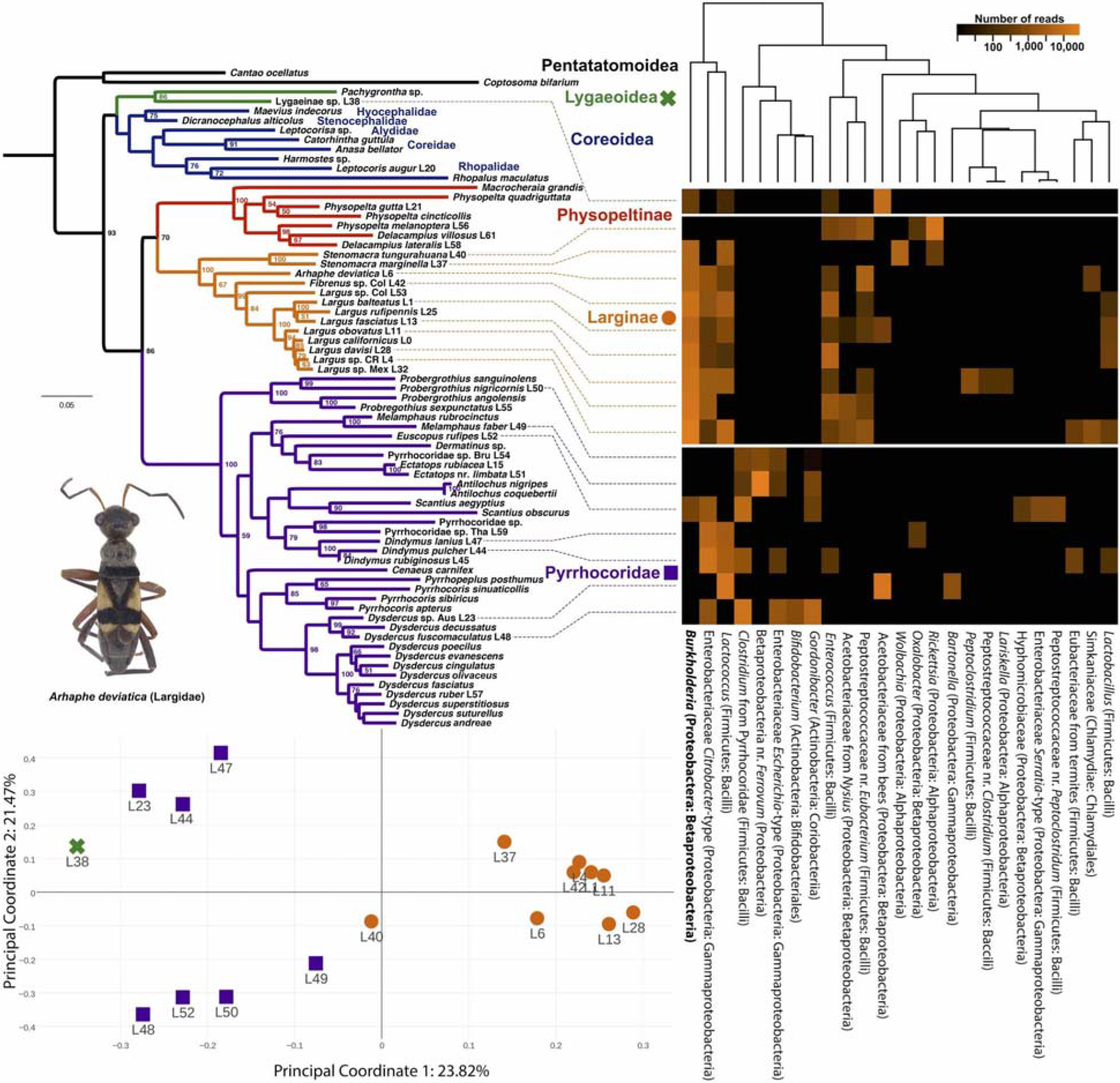
Maximum-likelihood phylogeny of Pyrrhocoroidea with bootstraps >50% displayed on branches and number of nucleotide changes indicated with scale bar indicating branch length corresponding to a mean of .05 nucleotide substitutions per site. Pentatomoidea was constrained as the outgroup. Displayed to the right is a heat map of bacterial OTUs from specimens indicated on the phylogeny with brightness corresponding to the log_10_ of number of Illumina reads divided by 10 or the log_10_ of reads after subtracting 1. The dendrogram on top of the heatmap represents clustering of OTUs based on their shared presence in samples. A principal coordinate analysis (PCoA) plot of abundance-weighted Unifrac distances is displayed on the bottom left.

### Bacterial associates of Pyrrhocoroidea

After quality control, we recovered a total of 319,021 paired-end Illumina reads (average of 18,766 per sample). We found a highly prevalent association of *Burkholderia* in gut extracts of all Largidae except for one specimen of *Stenomacra tungurahuana* (L40). We also recovered a previously described Pyrrhocoridae-associated *Clostridium* strain exclusively in each of our Pyrrhocoridae samples, although at a below 0.5% level for a *Dysdercus* species from Australia L23 and *Dindymus lanius* L47 (OTUs 6, 10, 19, 23 in Table S3; File S1). The presence of *Gordonibacter sp*. (OTUs 11, 28) was also observed exclusively in Pyrrhocoridae except for *Dindymus lanius* and only represented by two reads out of nearly 20,000 in *Di. pulcher* (below 0.5% of sample reads for *Probergrothius* and the *Dysdercus* species from Australia). We observed OTUs assigned to *Coriobacterium* only in our two sampled *Dysdercus* species at a sub 0.5% level (OTU 64 in Table S3).

The principal coordinate analysis plot displayed in Fig. 2 is that of a subsampled distance matrix based on abundance-weighted UniFrac distances (64) of OTUs and explains >45% of the variance in the data with the two plotted axes. Principal coordinate analysis of bacterial communities based on other distance measurements results in separate clusters of Largidae, Pyrrhocoridae and Lygaeinae in all but Bray-Curtis community dissimilarity matrices in which *Stenomacra tungurahuana* (L40) and *Stenomacra marginella* (L37) cluster separately from other Largidae (Fig S2.) near the Lygaeinae and Pyrrhocoridae samples. Two *Dindymus* species often cluster separately from most herbivorous Pyrrhocoridae sometimes also along with *Dysdercus* sp. L23 (Fig. 2; Fig. S2).

### Phylogenetics of *Burkholderia* associates

A phylogeny of *Burkholderia* symbionts based on full length 16S sequences (when available) was not concordant with host phylogenies (Fig. 3; top). There was no evidence of codiversification (Parafit: p-value = 0.5129). Five species of Largidae from geographically distant areas (Florida, Colombia, Argentina, Mexico and Costa Rica) harbored the same or very similar strains of *Burkholderia* which closely matched 16S sequences of PBE clade *Burkholderia*, including those of curated elite commercial inoculants used in agriculture for nodulation of legume crops in Brazil, housed at the SEMIA *Rhizobium* Culture Collection (e.g., 100% identical to *Burkholderia* sp. SEMIA 6385 / SEMIA 6382 isolated from *Piptadenia gonoacantha* / *Mimosa caesalpiniifolia* roots, FJ025136.1 / AY904775.1 [72]).

**Figure 3.**
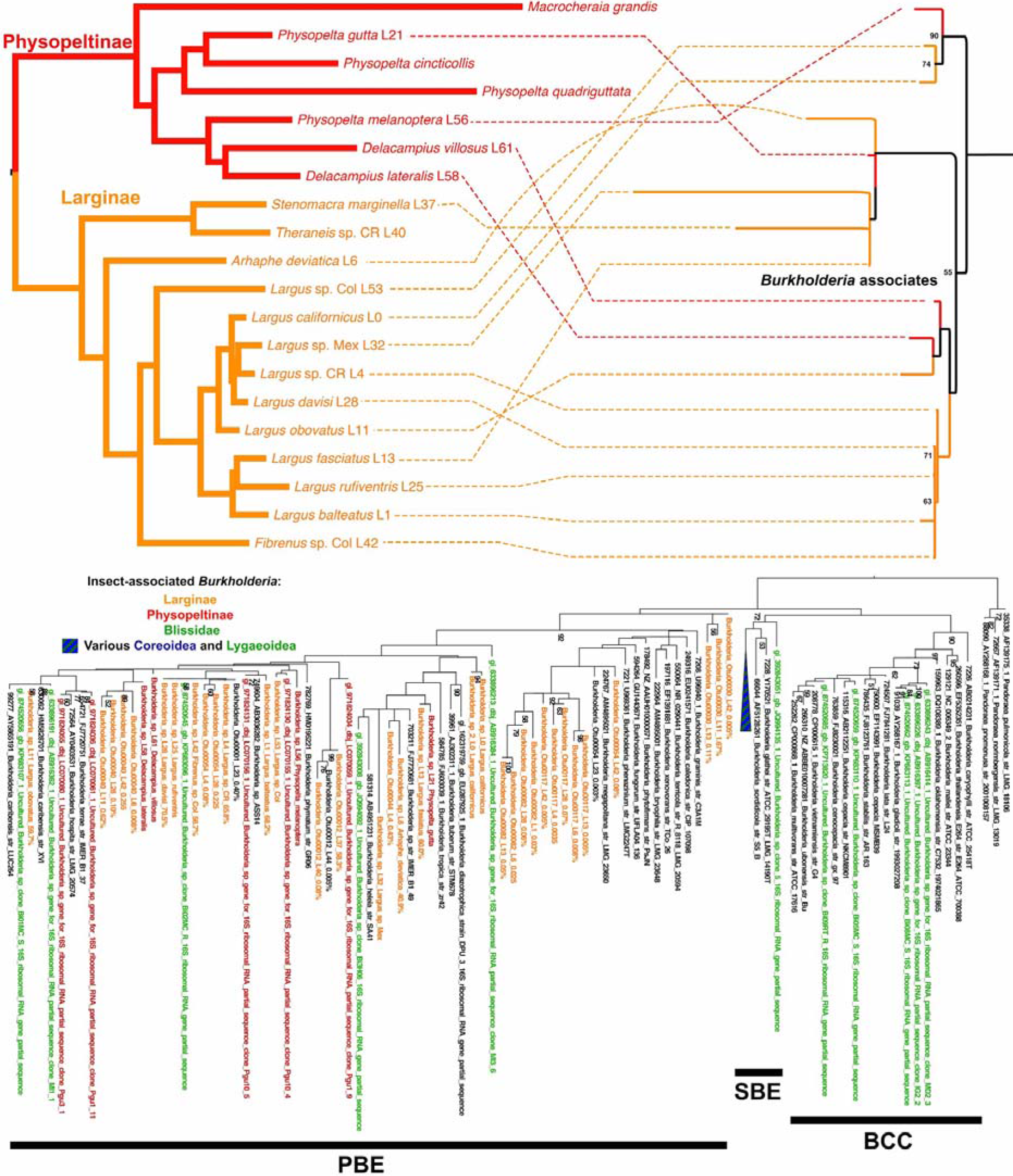
Bootstrap support >50% are displayed on bacterial phylogenies. Top. Maximally concordant host (left) and symbiont (right) maximum likelihood phylogenies as produced by TreeMap3 with linkages shown. Bottom. A phylogeny of newly obtained *Burkholderia* sequences with others from Genbank. Sequences are colored by host as in Figure 1. OTUs from the Illumina dataset (as well as full length 16S sequences from dominant *Burkholderia* strains) are followed by the percentage of reads they made up out of the total for that sample. One node consisting only of lygaeoid and coreoid associated *Burkholderia* in the SBE clade was collapsed for easier visualization.

A phylogeny constructed with all newly sequenced *Burkholderia* (orange and a subset of those in red) and relatives present on GenBank shows that all Largidae-associated strains of *Burkholderia* belong to the described PBE group (Fig. 3). Insect associates are closely related to species that are known nodulating bacteria of legumes and particularly of Mimosoideae (*Burkholderia diazotrophica, B. caribensis, B. phymatum, B. tuberum*; [73]) or other plant-associated nitrogen fixing species (*B. tropica, B. heleia*; [74]). All other non-Largidae heteropteran-associated *Burkholderia* fall within the SBE clade with the exception of a number of symbiont strains isolated from Blissidae (marked in green on Fig. 3). A tree with more comprehensive sampling of *Burkholderia* representatives including environmental isolates is shown in Fig. S3.

## Discussion

### Associated bacterial communities of Pyrrhocoroidea

We confirm an association of Largidae species from at least six genera with *Burkholderia* strains, specifically strains from plant-associated PBE clade, though there was also a single exception to this pattern, sample L40. Sample L40 (*Stenomacra tungurahuana*) instead possessed a high number of reads from *Rickettsia*. The *Dysdercus* specimen, L23, also had an unusual bacterial community profile with a high concentration of a species of Acetobacteraceae previously characterized from bees and also possessing a significant number reads from the intracellular pathogen, *Bartonella*. Both samples may be unrepresentative due to infection with another bacterium or could represent samples that suffered from decomposition or degradation of DNA after inadequate preservation in ethanol. Ethanol preservation has been shown to have an effect on sequenced bacterial diversity in other insects (75, 76). However, even these two samples cluster with other related members in most PCoA plots based on distance matrices except for that of Bray-Curtis distances. While L40 completely lacks any *Burkholderia* OTU, the *Burkholderia* strain present in sample L37 has less than 97% identity to 16S rRNA sequences from other largid-associated *Burkholderia* strains, thus it is not surprising that this similarity in genus composition would not be represented by phylogenetically-independent distance metrics. Members of two largid genera that we were not able to sample, *Macrocheraia grandis* and *Iphita limbata*, have been shown to possess caeca filled with rod-shaped bacteria potentially consistent with their identity as *Burkholderia (49, 50)*, though after culturing, isolates displayed differences with *Burkholderia*, such as being Gram-positive or spore-forming.

Although it was not our primary goal, we recovered a slightly more restricted distribution of some previously described Pyrrhocoridae symbionts than has been previously described. The most notable difference is the presence of *Coriobacterium* only in the *Dysdercus* species in our sampled representatives. Previously, *Coriobacterium* has been shown to make up high proportions of bacterial communities in members of the genera *Scantius, Pyrrhocoris* and *Dysdercus* (only the latter sampled in the current study) but represented <5% of reads also in *Antilochus, Probergothius* and *Dindymus*. Similarly, *Gordonibacter* has been shown to be present at low levels in many genera of Pyrrhocoridae including *Dindymus*, whereas we retrieved either extremely low or no numbers of *Gordonibacter* reads present in the representatives of *Dindymus* we sampled. In both of these predatory representatives, we instead found a microbiome dominated by a *Citrobacter*-type Enterobacteriaceae and a *Lactococcus* strain both of which appear to be common insect gut inhabitants in Largidae and perhaps other insects. The retention of any strict association of a predator with a bacterium, which we observe with the Pyrrhocoridae-associated *Clostridium* species is surprising as the evolution of a predatory life strategy in other Pentatomomorpha (such as the Asopinae [Pentatomidae] and Geocorinae [Geocoridae]) seems to negate dependence on any particular bacterium. This may reflect a relatively recent evolution of this predatory trophic strategy. The observation of well-developed caeca in one female specimen (L54) of Pyrrhocoridae should be investigated with fresh or acetone preserved specimens, if possible. Although vestigial caeca have been noted in *Dysdercus, Antilochus, Probergothius* and *Pyrrhocoris (51)*, it may be possible that at least some members of the well supported clade containing *Melamphaus* + *Dermatinus* + *Ectatops* + *Euscopus* retain functional caeca.

### Transmission method of *Burkholderia*

Although it has not yet been shown experimentally, all available evidence supports environmental reacquisition of *Burkholderia* by new generations of species of Largidae. The lack of evidence of *Burkholderia* in or on eggs or in lab-reared first instar nymphs as well as the lack of any sort of cophylogenetic signal suggest horizontal transmission. Booth (77) noted that nearly all first instar nymphs of *Largus californicus* reared in a laboratory setting died before molting, though eggs reared in the field and field-caught first instars brought to the lab were both viable to adulthood. This suggests that the first instar stage is when symbionts are acquired from the environment unlike in Alydidae, where symbionts are acquired during the second instar (26). Also suggestive of a horizontal transmission strategy is that in all other known cases of Heteroptera associated with *Burkholderia*, the symbiosis is overwhelmingly acquired from the environment (although with up to 30% vertical transmission in Blissidae; [(27)]).

Environmentally-acquired obligate symbiosis with bacteria is common in some microhabitats such as in soil or the ocean where it occurs in partnerships between nitrogen-fixing bacteria and legumes or bioluminescent bacteria and squids (78). However, this method of transmission for obligate symbionts is very rare in insects and association with *Burkholderia* in Heteroptera may be one of the oldest stably maintained symbiosis in insects, as the node containing all *Burkholderia*-associated members has been dated to the early Cretaceous era ~130 mya (54). Such associations have the potential to evolve towards pathogenicity of symbionts as horizontal acquisition can select for pathogenic or cheating phenotypes (79) and other similar systems show policing by hosts to punish symbionts which do not participate (80). The gut crypts of Largidae and other heteropterans may provide a much-needed mechanism to police symbionts as individuals crypts could be modulated individually, though this has not yet been shown.

### Evolutionary transitions of symbiont complexes

The exclusive association of Largidae with PBE clade *Burkholderia* unlike all other *Burkholderia*-associated Heteroptera is unlikely to be incidental. The Pyrrhocoroidea have now been shown to be one of two lineages of *Burkholderia*-associated Heteroptera, with the Largidae the earliest diverging single family which still retains such a symbiosis. It is likely that the ancestor of all *Burkholderia*-associated Heteroptera was also associated with *Burkholderia* but it is not as clear which clade of *Burkholderia* or if there was any specificity at all. Perhaps this ancestor had the same lack of specificity towards *Burkholderia* that members of Blissidae seem to display when associating with different strains of *Burkholderia* spanning the three described clades. As lineages diverged, different mechanisms of specificity may have evolved that selected for particular subclades of *Burkholderia* with increased evolutionary benefits for that lineage depending on its biology. Many members of the PBE clade of *Burkholderia* are rhizobia that form nodules on legumes and these symbiotic properties tend to be tied to symbiotic genomic islands that contain genes for nodulation and nitrogen fixation that are easily horizontally transferred (81). Other PBE *Burkholderia* are also nitrogen-fixing and associated with plants but their genomic architecture is not yet known (74, 82). Nitrogen fixation may be the role of these symbiotic bacteria to their largid hosts as has been suggested of other insect symbionts from lineages of bacteria that undergo nitrogen-fixing symbioses with plants, such as in herbivorous ants (83). It is possible that an increased efficiency of nitrogen fixation (e.g., of elite commercial inoculants) is the mechanism that selects for similar strains in geographically disparate Largidae. These bacteria can presumably live within either of two symbiotic systems or as a free-living bacterium within the environment and must face different selective pressures in each of these states.

Strict vertical transmission of symbionts, especially intracellular symbionts, can lead to rapid degradation of symbiont genomes and may be non-beneficial in some respects, trapping hosts and symbiont in “an evolutionary rabbit hole” (6). However, the ancestor of Heteroptera achieved independence from intracellular symbionts, but after re-evolving herbivory, some members developed new associations with extracellular bacteria. In the Pentatomoidea, these symbionts are primarily vertically transmitted, although there is accruing evidence suggesting that vertically transmitted symbionts can be supplanted by environmental bacteria, thus replacing any possibly degraded symbiont genome (84). For *Burkholderia*-associated members, symbionts are primarily acquired from the environment every generation thus avoiding any transmission bottleneck that could lead to genome degradation (22). However, within the Pyrrhocoridae, there has been a transition to vertically transmitted extracellular symbionts, and in some lineages of Lygaeoidea, several reversions to vertically-transmitted intracellular symbioses, the ancestral condition of non-heteropteran Hemiptera (Fig. 4). It has been suggested or implied that these transitions to vertically transmitted symbionts has been driven by evolutionary pressures tied to host plant specificity (20, 51). But host plant specialists are also widespread in species that retain environmentally-acquired symbionts (13, 85). Perhaps, there are more specific evolutionary pressures that lead to this more stable but eventually degenerate form of symbiosis.

**Figure 4.**
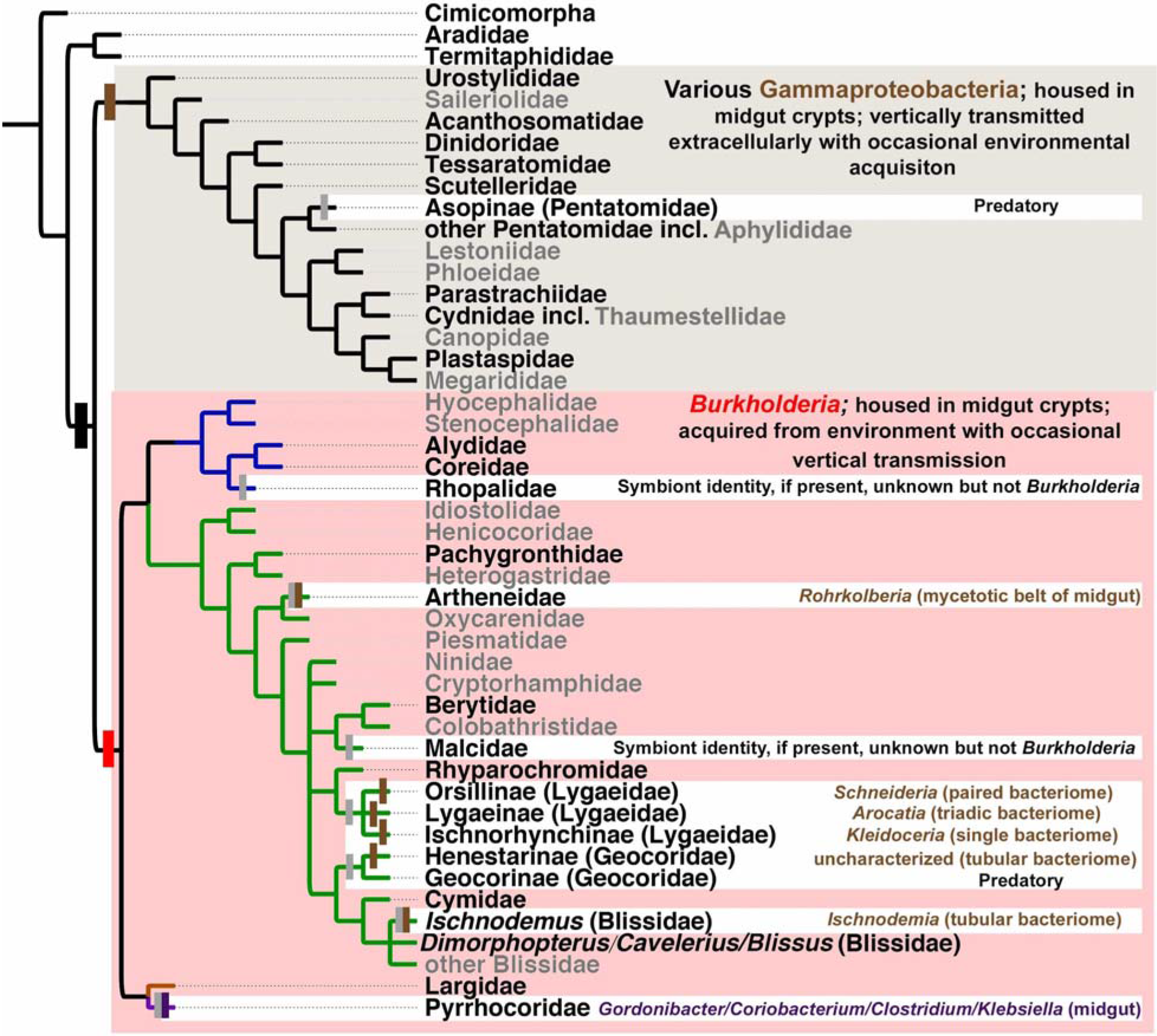
The large black mark indicates the evolution of caeca. Grey marks indicate a loss of these caeca (vestigial in Pyrrhocoridae). Brown marks indicate the evolution of association with gammaproteobacteria, red with *Burkholderia*, and purple with a consortium of bacteria including two Actinobacteria. Taxa in grey have not yet been examined. The phylogeny is based on relationships recovered in this paper (for Coreoidea) and well-supported relationships from Bayesian and likelihood analyses based on six Hox gene fragments (87). Families not represented in that analysis or with not well-supported relationships were placed using successive weighting parsimony analyses based on morphology as in (57) for Lygaeoidea and (88) for Pentatomoidea. References for symbiotic associates are as in text (20–22, 32). Only one member of Lygaeinae (*Arocatus longiceps*) has so far been demonstrated to be associated with a symbiont (20). Oxycarenidae are purported to possess an unpaired bacteriome but the identity of the bacterial inhabitant is unknown (89).

There remains outstanding questions with respect to the mechanism and evolutionary benefit of specificity to different clades of *Burkholderia* in different heteropterans. Generalist vs. specialist evolutionary pressures may play a role. Many families remain to be characterized (Fig. 4) especially using modern molecular methods and other families demonstrate such a diversity of symbiotic complexes in known members that many more members should be examined. Particularly, the identity of any symbiont present in members of the two small families sister to all other Coreoidea, Hyocephalidae and Stenocephalidae (the former, associated with *Acacia* and *Eucalyptus* seeds, and the latter, specialists on seeds of Euphorbiaceae [42]), may contribute to our understanding of how the evolution of strict associations with clade-specific *Burkholderia* arose. While the genome of a SBE clade *Burkholderia* isolated from Alydidae has been published (86), it has so far not provided much insight into the probable function of this symbiont. Unlike strict symbionts, these bacteria have large genomes and thus it is more difficult to discern function from the retention of gene sequences alone. The sequencing of PBE clade *Burkholderia* symbionts from Largidae, which in at least one case, is easily cultured, may provide some insight into shared genes between these symbiotic strains as well as differences among them.

## Acknowledgments

We would like to thank Kaleigh Russell for assistance in culturing bacteria, Alex Knyshov for guidance with FISH microscopy and Paul Masonick for help with collecting *Largus californicus*.

## Funding information

This study was funded by the Dr. Mir S. Mulla and Lelia Mulla Endowed Scholarship Fund and the National Science Foundation Graduate Research Fellowship awarded to E. R. L. Gordon and initial complement funds awarded to to Q. S. McFrederick by University of California, Riverside.

## Supplementary data

Table S1. Specimen information and accession numbers.

Table S2. PCR primers and conditions

Table S3. OTU count table

File S1. Representative OTU fasta file

Figure S1. Phylogeny of Pyrrhocoroidea with all bootstrap supports

Figure S2. PCoA plots based on other distance metrics.

Figure S3. Untrimmed *Burkholderia* phylogeny

